# OpenFIBSEM: an application programming interface for easy FIB/SEM automation

**DOI:** 10.1101/2022.11.01.514681

**Authors:** Patrick Cleeve, David Dierickx, Genevieve Buckley, Sergey Gorelick, Lucile Naegele, Lachlan Burne, James C Whisstock, Alex de Marco

**Affiliations:** Department of Biochemistry and Molecular Biology, Biomedicine Discovery Institute, Monash University, 3800 Clayton Victoria, Australia; Ramaciotti Centre for cryo-Electron Microscopy, Monash University, 3800 Clayton Victoria, Australia; Monash Centre for Electron Microscopy, Monash University, 3800 Clayton Victoria, Australia; EMBL Australia, Monash University, 3800 Clayton Victoria, Australia

**Keywords:** Focused Ion Beam microscopy, automation, Python, API, Microscopy, controller

## Abstract

Automation in microscopy is the key to success in long and complex experiments. Most microscopy manufacturers provide Application Programming Interfaces (API) to enable communication between a user-defined program and the hardware. Although APIs effectively allow the development of complex routines involving hardware control, the developers need to build the applications from basic commands. Here we present a Software Development Kit (SDK) for easy control of Focussed Ion Beam Scanning Electron Microscopes (FIB/SEM) microscopes. The SDK, which we named OpenFIBSEM consists of a suite of building blocks for easy control that simplify the development of complex automated workflows.

## Introduction

Focused Ion Beam Scanning Electron Microscopes (FIB/SEM) have been widely used across multiple fields: from semiconductors to materials and life sciences(Lepinay and Lorut, 2013; Phaneuf, 1999; Schaffer and Wagner, 2008; Villinger et al., 2012). The most common uses for FIB/SEMs are sample preparation for transmission electron microscopy (TEM), commonly known as lamella preparation (Rigort et al., 2012; Schaffer et al., 2012); FIB lithography (binary, grayscale, or additive) (Garg et al., 2019; Gorelick et al., 2019; Gorelick and De Marco, 2018), and three-dimensional imaging (Slice&View or metrology) (Kizilyaprak et al., 2014; Mammadi et al., 2020; Villinger et al., 2012). Regardless of the application, automation, customisation of the workflow, and real-time data analyses are critical. The effects of automation regarding throughput and success rate have been amply demonstrated on multiple occasions in microscopy. For example, in TEM imaging, the introduction of SerialEM, Leginon, and other open-access TEM control software (Carragher et al., 2000; de la Cruz et al., 2019; Potter et al., 1999; Suloway et al., 2009; Zheng et al., 2007) allowed high-throughput single particle and tomographic acquisition, as well as the introduction of complex and non-conventional acquisition methods such as the dose-symmetric tilt-series (Hagen et al., 2017). In light microscopy, Micro-manager (Edelstein et al., 2010; Edelstein et al., 2014) allowed easy control of commercial and custom-built setups and the development of automated procedures for complex data acquisition (Pitrone et al., 2013). Both the abovementioned cases effectively represent an easy interpreter of the microscope and detector manufacturers with both a graphical and a scriptable user interface.

Microscope manufacturers provide Application Programming Interfaces (API) to allow communication with the instrumentation. The classes in the APIs comprise basic commands and calls to low-level actions such as stage movements (absolute or relative) or basic image acquisition. In this work, we focus on the ThermoFisher API Autoscript. The library grants access to all the core functionalities of the microscopes, but it does not provide significantly advanced functionality. For example, it is easy to define image acquisition using Autoscript, but it is not possible to customise the autofocus routine with arbitrary parameters.

Currently, there are multiple commercial products available to run automated procedures. For example, AutoSlice&View allows the guided setup of FIB/SEM tomography data acquisition. The program has an easy-to-follow graphical user interface and allows for a certain degree of flexibility in the setup. Still, due to how data is stored in the memory, it shows significant lag on long acquisitions, and it lacks the possibility to automatically re-align the Field of View (FOV) based on the image content. The main consequence of this limitation is that upon acquisition of large volumes, the FOV of interest moves out of the imaged region once a significant depth has been milled. The abovementioned example highlights how easy it is for a research-oriented commercial product not to anticipate a use case.

In most cases, the manufacturers will address these limitations, but between the user feedback and the implementation of the desired feature, up to two years can easily pass. Further, the commission may never occur for some features deemed minor or niche. As such, due to the inability to alter the program flow or customise how lower-level operations are performed, researchers are often unable to pursue a research direction due to the cost and complexity of redeveloping solutions.

There are at least two examples in TEM imaging where open-source software was developed and successfully pushed the development of custom complex workflows: SerialEM (Mastronarde, 2005) and Leginon (Carragher et al., 2000). Here, both packages are built on the API provided by the microscope manufacturers but provide significantly advanced functionality and, most importantly, the possibility to easily script and create novel pipelines. For example, the recent development of py-EM led to the possibility controlling TEM microscopes using Python (Schorb et al., 2019). This high level of flexibility associated with the increased accessibility given by the library led to the development of other packages such as PACEtomo (Eisenstein F et al., 2022). Finally, both, SerialEM and Leginon extend their support beyond a single manufacturer, making it easier for a novice user to learn the control of multiple instruments sharing a common interface.

Here we introduce OpenFIBSEM, a python-based Software Development Kit (SDK) that builds on the manufacturer’s API, providing a simplified way of interacting with the microscope and designing custom workflows. The package comprises a collection of classes that aid imaging, stage movement, manipulator control, system calibration and easy integration with a graphical user interface. Currently, OpenFIBSEM only supports ThermoFisher instruments, but we plan to extend this to other manufacturers in the future to enable users to develop automated routines on multiple instruments with a stable syntax; therefore, simplifying the adoption of complex workflows across facilities and instruments.

## Results and Discussion

OpenFIBSEM aims to enable fast workflow development while allowing the low-level customisation required by researchers (Figure 1). The library is designed to provide high-level abstractions of common operations, such as image acquisition and movement, and reusable components, such as automated calibration procedures. In addition, the library includes several reusable user interface components (PyQt / Napari) for common user operations (e.g. movement). Ideally, the library should abstract microscope operation from the underlying hardware, allowing the user to focus on what they want to do rather than how to do it.

**Figure 1:**
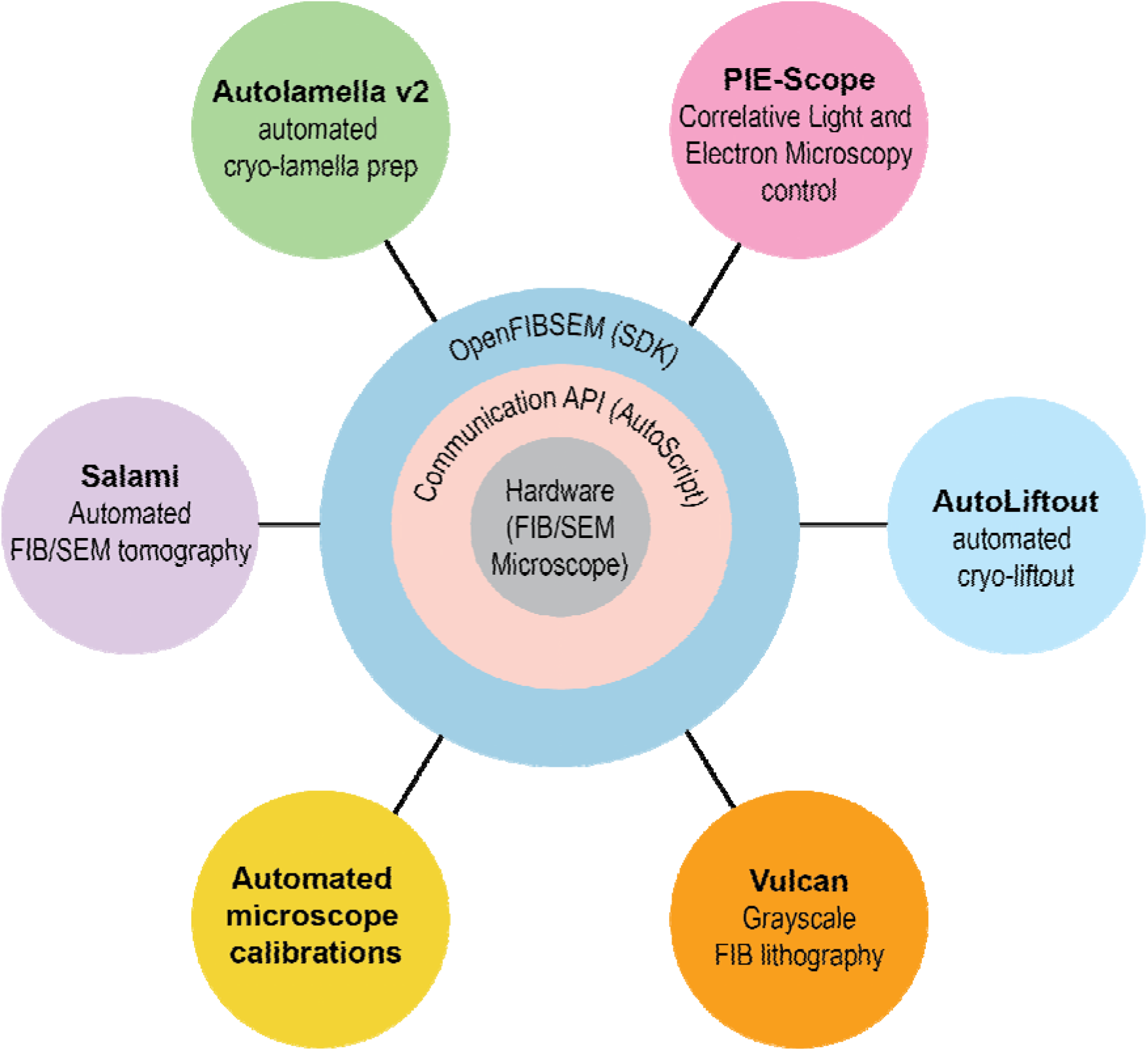
The architecture concept. The diagram shows the lowest level and therefore furthest away from the user is the hardware. The communication API provides access and control of the hardware, the Backend SDK provides a user-friendly control therefore enabling an easy development of user-specific applications. The outermost ring provides a list of examples of the user-specific applications that can easily be developed when a high-level API is present.

The library can be downloaded from www.github.com/demarcolab/fibsem and is organised according to the following structure:

### Core

- **Acquire**: A collection of helper functions for the simple definition of imaging settings enabling image acquisition and post-processing.
- **Milling**: A collection of helper functions to set up and run focused ion beam milling routines. This collection includes some patterning examples.
- **Movement**: Functionality for the stage and needle movement, corrected for both the view perspective and stage orientation.
- **Structures**: A collection of classes defining the data structures for a correct interpretation of the settings and types.
- **Utils**: A collection of general microscope utilities, including checks for network connection and filesystem access.

### Automation

- **Alignment**: Provides the tools for automated alignments of the sample position to correct for drift or to correct the eucentric position.
- **Calibration**: Includes multiple automated calibration routines and helps manage the microscope state.
- **Validation**: A collection of automated validation functions to check and set the desired microscope settings. The user can configure all the settings.

### Image Analysis

- **Imaging**: A collection of helper functions for simple image analysis and manipulation, including filtering and masking.
- **Detection**: Comprises functions for the automated detection of common features present in conventional workflows, such as the manipulator (NeedleTip class) or the post of a half-moon grid (LandingPost class).
- **Segmentation**: A library providing tools for implementing a deep learning-based segmentation workflow, including labelling, training and inference.

### Utilities

- **Conversions**: Provides standard conversions between coordinate systems used in the microscope and allows converting distances in images between pixels and meters.
- **UI**: A collection of basic user interface windows in PyQt designed for basic interaction with OpenFIBSEM. Two examples are the movement window and the detection window.

#### Getting started

The main goal of the package is to simplify the performance of general tasks within the microscope capabilities to enable rapid development. The library requires a general familiarity with Python programming; however, the setup process is concise. The example below shows how to connect to the microscope, define the imaging parameters and acquire an image using the electron beam with the automated contrast selection.

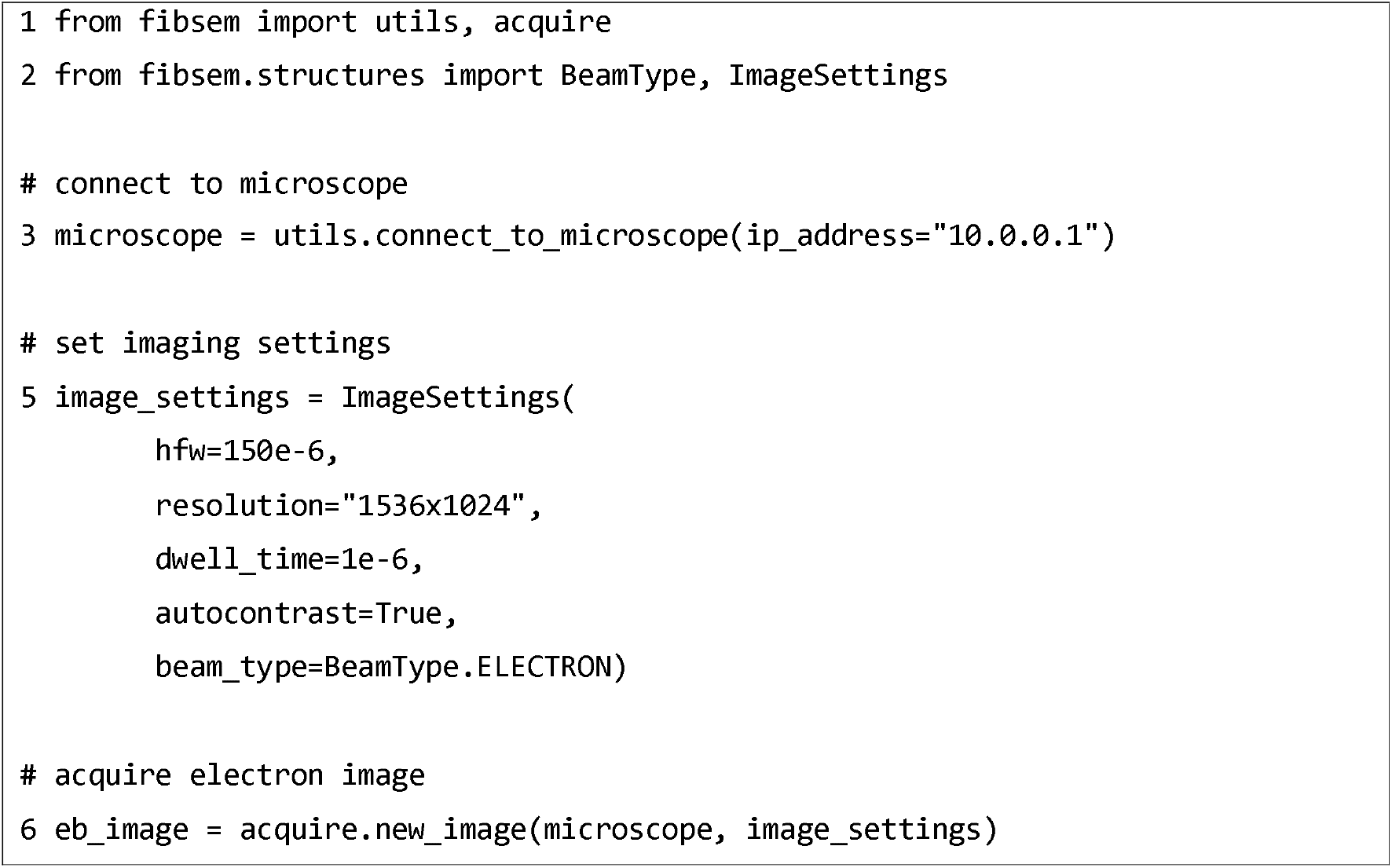

Although acquiring an image is a simple case, it is the most commonly performed microscope operation. By analysing the difference between the code required using OpenFIBSEM and the one directly implemented from AutoScript, it is immediately evident that the complexity has decreased and it is now possible to obtain the same result without the need of traversing into three separate modules (imaging, beams, and auto_functions).

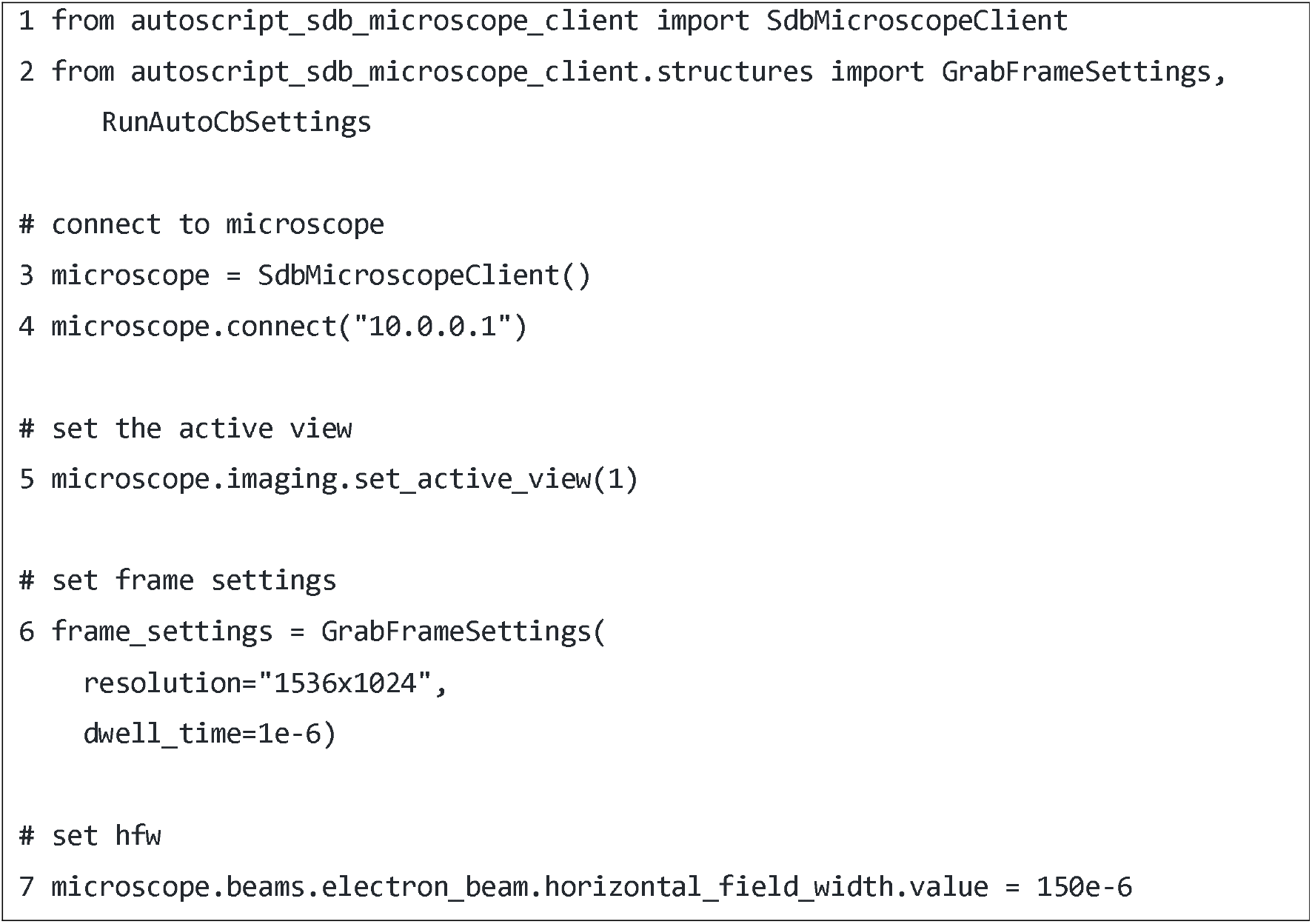
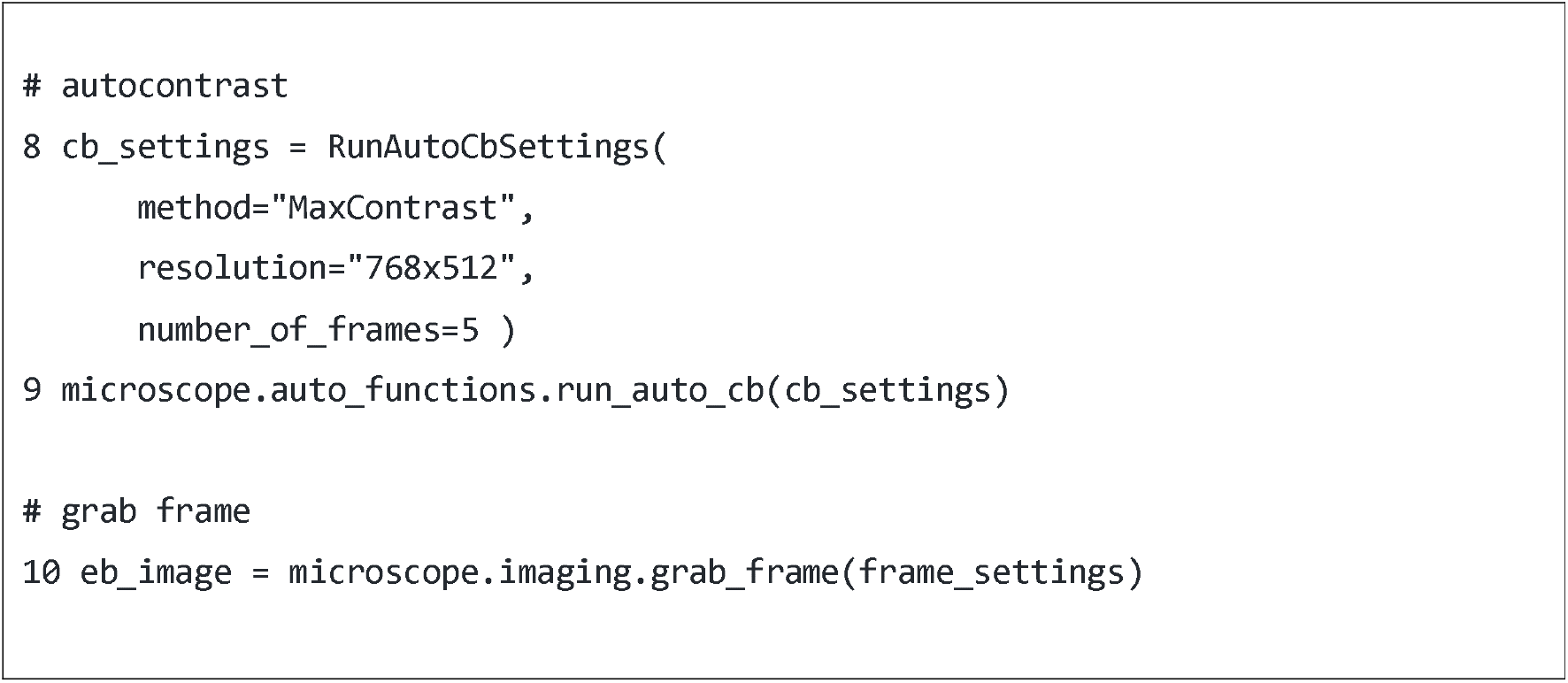

#### Automating routines

One of the most common actions performed when using a FIB/SEM is focusing; this is crucial not only to obtain optimal images but also to provide a measurement of the effective position of the sample along the electron axis. Accordingly, in automated procedures, a robust autofocus routine is critical.

The autofocus routine is an excellent example of an automated procedure that is available but inefficient. Autoscript provides a standard autofocus routine that uses the full image and focusses with a standard interval (+/-10μm). In many cases, this routine is not flexible enough; for example, if the sample is highly dose-sensitive, using a sub-region is critical to avoid damage because of repeated exposure. Further, re-focusing is likely required if the stage movements are significant and the sample is irregular. In this case, the standard procedure often fails because the interval for the autofocus may not be large enough. Below is an example of a simple routine developed with OpenFIBSEM enabling custom autofocus.

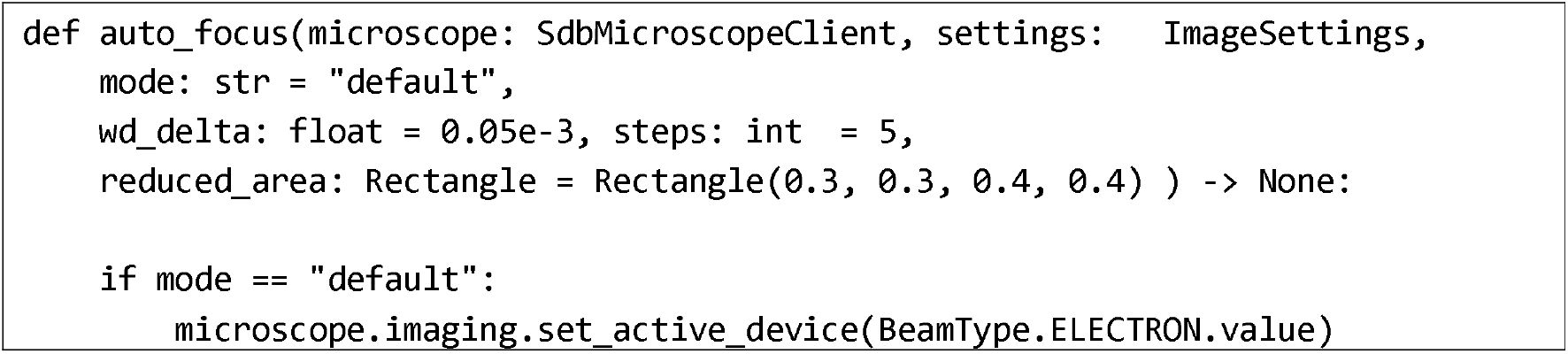
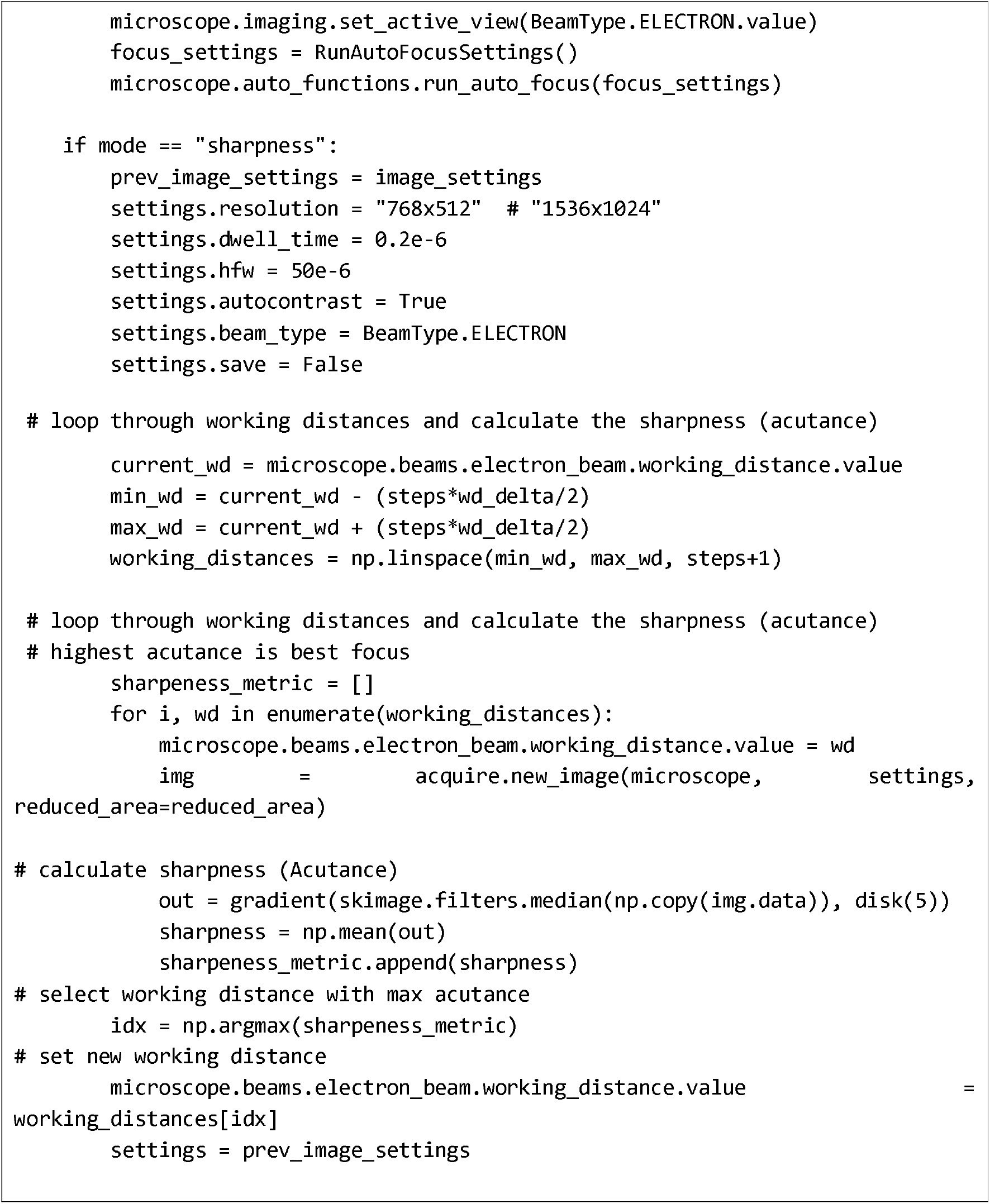

#### Enhanced Functionality

The AutoScript API provides functionality to move the stage in two coordinate systems (relative and raw). The raw coordinate system is based on the physical cartesian coordinates, while the relative coordinate system is based on the relative distance from the beam (defined by the focus and link routine). When moving using AutoScript or the xT-UI, the stage will move correctly and maintain eucentricity between both the Electron and Ion beams. However, if the stage shuttle has a pre-tilted angle (such as for lamella preparation to provide additional access for the Ion beam), neither the API nor the UI will move correctly and degrade eucentricity over time as there is no way to tell the microscope that the stage is pre-tilted.

OpenFIBSEM provides corrected movement functions for both the stage and manipulator. The user can provide the pre-tilt of the shuttle in the settings, and the library will calculate the corrected movements to maintain eucentricity. Moreover, the user doesn’t need to spend time understanding the implementation details regarding different coordinate systems, stage movements, and hardware and, therefore, can focus on research. Figure 2 shows the difference between baseline and corrected movements for a stage with a 35-degree pre-tilt angle. The user clicks the same position in xT-UI, and openFIBSEM and the actual positions after movement for each beam are shown.

**Figure 2:**
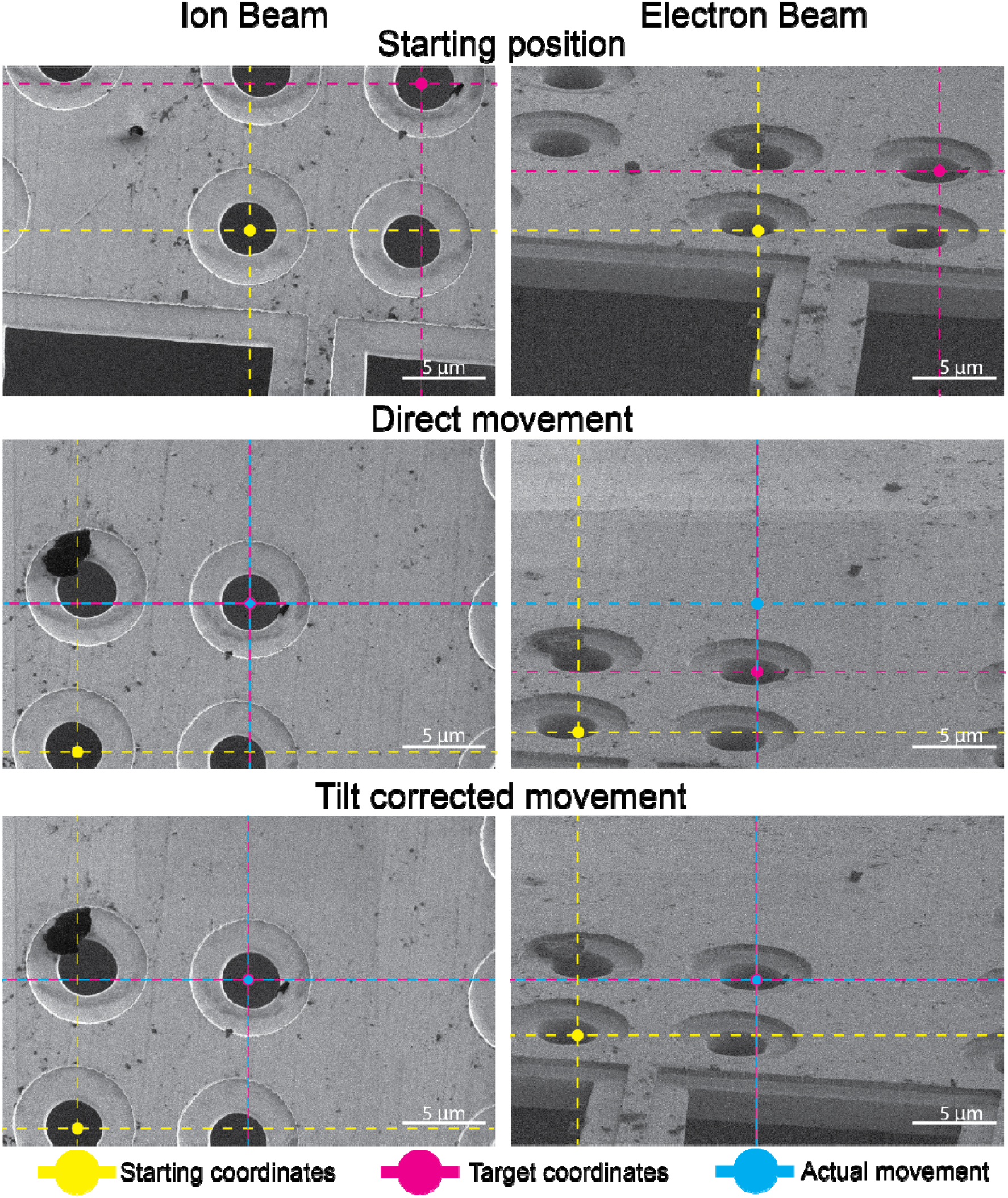
OpenFIBSEM Corrected Movement. Given the orientation of the beams relative to the sample the XY coordinate axes of the stage correspond to those of the Electron Beam view. Accordingly, as visible from the left column it is shown how the selection of a destination drives the correct movement of the stage to the target location. From the Ion Beam view a direct movement based on the distance between the starting point and the desired location provides an incorrect movement. This mismatch between the target and the actual reached position depends on the fact that there is a difference between the coordinate system of the stage and the beam. Further, if the sample is mounted with a tilt relative to the XY plane of the stage this mismatch becomes particularly evident from the Ion Beam view and must be accounted for. The last panel shows the effect that tilt correction has when calculating the correct shift.

#### Common Workflow Tools

OpenFIBSEM provides a set of common tools enabling multiple workflows utilised in imaging. For example, deep learning has been successfully implemented for denoising and segmentation (a couple of examples repositories can be found here: https://github.com/HenriquesLab/ZeroCostDL4Mic, https://github.com/juglab/DenoiSeg). However, specific implementations often don’t provide the tools or flexibility for researchers to utilise models or only cover a specific aspect of deep learning (usually training the model). This excludes the most time-consuming and technically difficult aspects of using a deep learning model: labelling and deployment. To overcome this problem, here we include a workflow for collecting microscopy images, labelling them, training a segmentation model, and deploying it in a workflow.

We have created a labelling user interface using napari (Sofroniew et al., 2022) and Dask (Rocklin, 2015), allowing a user to label hundreds of images efficiently. The user can then train and test their choice of prebuilt models from segmentation-models-Pytorch (Paszke et al., 2019) by editing a simple configuration file (YAML). We also provide a simple wrapper class for deploying the model into an openFIBSEM workflow. An example of the proposed workflow can be found in figure 3, with example outputs of each stage of the pipeline.

**Figure 3:**
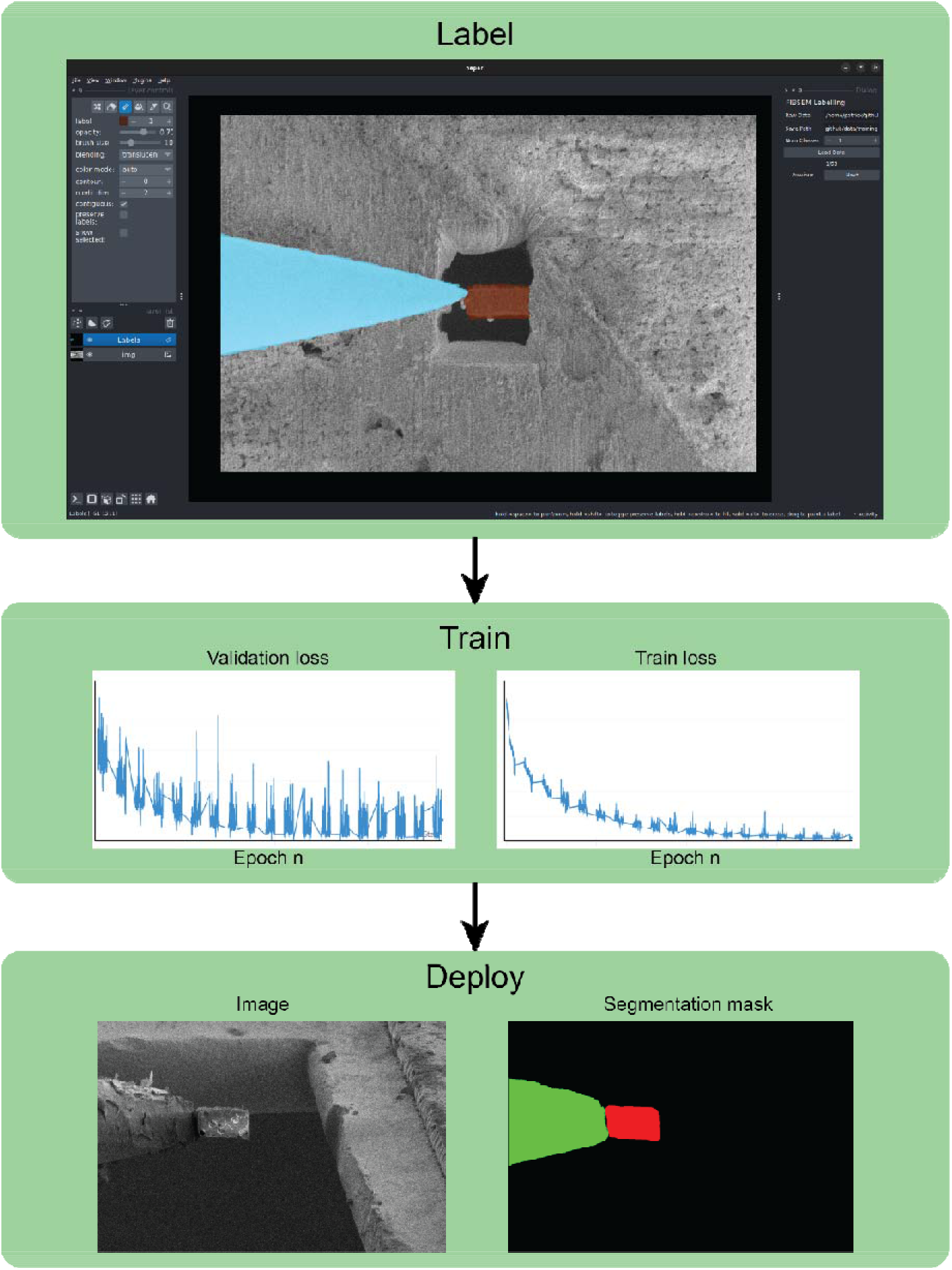
The U-Net assisted segmentation workflow integrated in OpenFIBSEM. The toolkit includes a user interface based on napari to import and label (segment) the images to train the model, a training setup and the code to easily deploy the segmentation model to identify correctly features of interest. The example in the figure shows the labelling and the recognition of the needle micromanipulator and a lamella.

#### User Interface

To enable non-expert users to utilise FIBSEM machines to pursue research, workflows often require image visualisation and user interaction. To this end, we developed a library of common user interface components and tools based on PyQt and Napari that can be used as standalone or as building blocks to be integrated into larger applications.

An example of a user interface provided with OpenFIBSEM, that can be used standalone or integrated into larger workflows and application, is the movement window. This application provides a user interface for users to double-click to move (mirroring the microscope user interface interaction). This applet allow navigation in two different movement modes, stable and eucentric. Stable movements (discussed above) consist of standard stage moves that maintain eucentricity between beams, while eucentric mode can correct the eucentricity between beams. The movements in both cases account for the sample pre-tilt and assume a flat sample, therefore in the case of a sample displaying large variations in height across the surface corrections will be required more often when moving. Over time, we aim to add additional components and improve the functionality and modularity of these interfaces.

#### Examples of workflows developed with OpenFIBSEM

To highlight the possibilities offered by the OpenFIBSEM package, we developed a few examples of automated routines covering multiple use cases (Figure 1). All the exemplary code can be found in the “Example” directory of the repository.

The most straightforward case use consists of performing grey-scale lithography using a pattern imported from images. This is common practice in microfabrication, and the example we present consists of fabricating the profile of a lens. The applet includes the interactive movement along the sample using the eucentric corrected movement and ensures that the image file format is compatible with the pattern definition. Patterns are created as 8-bit grayscale images and then converted to a 24-bit bitmap as required by the system.

Another case is the serial fabrication of lamellae on grids. This use-case is extremely popular in the life sciences, and the automation of this procedure has been presented in a couple of articles (Buckley et al., 2020; Klumpe et al., 2021).

The first example of this procedure was published by our group and was a complex package; with OpenFIBSEM, the core of the program can be scripted extremely quickly. A working example of this procedure can be found in the Example directory named: “autolamella.py”. Although this is only a simple example, it highlights the potential for new users and groups to develop their own automation routines.

The last example is the performance of FIB/SEM tomography. This is common practice across multiple disciplines, from the material to the life sciences. The workflow consists of imaging a surface with the SEM, polishing a defined amount of material from the surface using the FIB and imaging the newly exposed surface. This is then iterated until the desired volume has been acquired. The simplest setup does not require any stage movement, and the SEM imaging is performed at an angle using the dynamic focusing option to compensate for the focal gradient. In these conditions, the code is reduced to a loop where the sample surface is first imaged, and then the surface is polished with the FIB. A re-alignment step can be added in the loop to compensate for any drift. Among the examples available in the repository, we show a short version of the FIB/SEM tomography workflow developed in less than 40 lines of code using OpenFIBSEM.

Although these examples were intended as simple demonstrations or re-implementations of existing workflows to provide users with a set of templates to start using OpenFIBSEM, we immediately found use cases and users for these programs. Accordingly, these developed into standalone projects.

### Vulcan

Vulcan is a project for simplifying and enhancing grayscale lithography manufacturing using OpenFIBSEM. It contains tools for pattern conversion, milling calibration, and workflow tools. It can be used, for example, in conjunction with the optical simulation package Juno (https://github.com/DeMarcoLab/juno) to design, develop, manufacture and validate micro-optical elements. https://github.com/DeMarcoLab/vulcan

### Salami

Salami is a custom slice and view project to enable custom data collection strategies. The project is currently being used to screen and optimise data collection strategies for serial FIB/SEM tomography of large samples. The flexibility of the package allows for testing The iterative development for the project is shown in Figure 4, and the repository can be accessed from https://github.com/DeMarcoLab/salami.

**Figure 4:**
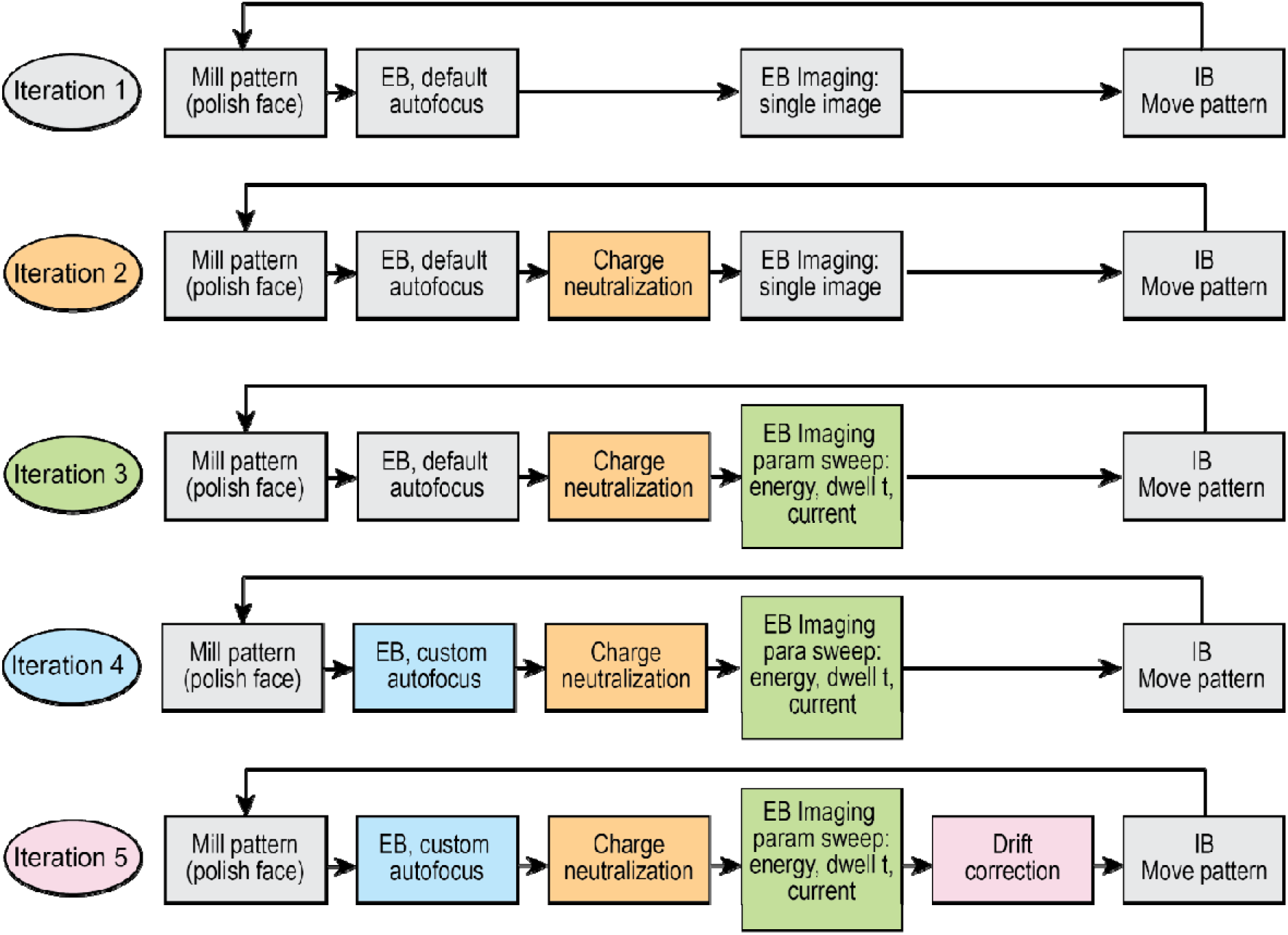
Diagram of an example of possible evolution of a protocol. Here we show 5 iterations during the development of a FIB/SEM tomography workflow. The colours are representative of the components that have been added at each iteration. The example shows how the initial iterative sequence of milling a surface, imaging, and repositioning of the milling pattern can be improved by adding a step to neutralise the charge left by the ion beam, then the imaging parameters can be changed, or screened. Finally, the autofocus routine can be improved by using a custom algorithm and finally a drift correction can be applied.

## Conclusions

In this article we introduced a high-level SDK enabling the quick and robust development of automated pipelines for FIB/SEM data acquisition and microfabrication. The increased level of abstraction provided by this package compared to the microscope API does not only make the development of new routines easier, but it is also integrated with numerous accessory functions that can be used to extend the capability of the microscope by integrating complex image processing pipelines.

The current version of OpenFIBSEM has been developed around the ThermoFisher Autoscript API. We inted to maintain this package, while extending the support to other manufacturers, with the goal of providing a common high-level programming language across multiple instruments. The code is freely available under the MIT licence agreement at www.github.com/demarcolab/fibsem. To request new features or report bugs users are asked to submit the requests through GitHub.

## Acknowledgements

PC was supported by the Australian Research Data Commons project XN006. DD, GB, JCW were supported by the Australian Research Council Laureate Fellowship FL180100019. SG and AdM were supported by the Human Frontiers Science Program grant RGP0017/2020.

## Authors contributions

PC conceived and developed the library architecture and wrote the article, DD, GB, SG LB, LN developed the library. AdM and JCW managed the project and wrote the article.

## References

Buckley, G., Gervinskas, G., Taveneau, C., Venugopal, H., Whisstock, J.C., and de Marco, A. (2020). Automated cryo-lamella preparation for high-throughput in-situ structural biology. J Struct Biol 210, 107488.

Carragher, B., Kisseberth, N., Kriegman, D., Milligan, R.A., Potter, C.S., Pulokas, J., and Reilein, A. (2000). Leginon: an automated system for acquisition of images from vitreous ice specimens. J Struct Biol 132, 33–45.

de la Cruz, M.J., Martynowycz, M.W., Hattne, J., and Gonen, T. (2019). MicroED data collection with SerialEM. Ultramicroscopy 201, 77–80.

Edelstein, A., Amodaj, N., Hoover, K., Vale, R., and Stuurman, N. (2010). Computer control of microscopes using microManager. Curr Protoc Mol Biol Chapter 14, Unit14 20.

Edelstein, A.D., Tsuchida, M.A., Amodaj, N., Pinkard, H., Vale, R.D., and Stuurman, N. (2014). Advanced methods of microscope control using muManager software. J Biol Methods 1.

Eisenstein F, Yanagisawa H, Kashihara H, Kikkawa M, Tsukita S, and Danev, R. (2022). Parallel cryo electron tomography on in situ lamellae. Biorxiv, 487557.

Garg, V., Chou, T., Liu, A., De Marco, A., Kamaliya, B., Qiu, S., Mote, R.G., and Fu, J. (2019). Weaving nanostructures with site-specific ion induced bidirectional bending. Nanoscale Adv 1, 3067–3077.

Gorelick, S., De Jonge, M.D., Kewish, C.M., and De Marco, A. (2019). Ultimate limitations in the performance of kinoform lenses for hard x-ray focusing. Optica 6, 790–793.

Gorelick, S., and De Marco, A. (2018). Fabrication of glass microlenses using focused Xe beam. Opt Express 26, 13647–13655.

Hagen, W.J.H., Wan, W., and Briggs, J.A.G. (2017). Implementation of a cryo-electron tomography tilt-scheme optimized for high resolution subtomogram averaging. J Struct Biol 197, 191–198.

Kizilyaprak, C., Bittermann, A.G., Daraspe, J., and Humbel, B.M. (2014). FIB-SEM tomography in biology. Methods Mol Biol 1117, 541–558.

Klumpe, S., Fung, H.K., Goetz, S.K., Zagoriy, I., Hampoelz, B., Zhang, X., Erdmann, P.S., Baumbach, J., Muller, C.W., Beck, M., et al. (2021). A modular platform for automated cryo-FIB workflows. Elife 10.

Lepinay, K., and Lorut, F. (2013). Three-dimensional semiconductor device investigation using focused ion beam and scanning electron microscopy imaging (FIB/SEM tomography). Microsc Microanal 19, 85–92.

Mammadi, Y., Joseph, A., Joulain, A., Bonneville, J., Tromas, C., Hedan, S., and Valle, V. (2020). Nanometric metrology by FIB-SEM-DIC measurements of strain field and fracture separation on composite metallic material. Mater Design 192.

Mastronarde, D.N. (2005). Automated electron microscope tomography using robust prediction of specimen movements. Journal of Structural Biology 152, 36–51.

Paszke, A., Gross, S., Massa, F., Lerer, A., Bradbury, J., Chanan, G., Killeen, T., Lin, Z.M., Gimelshein, N., Antiga, L., et al. (2019). PyTorch: An Imperative Style, High-Performance Deep Learning Library. Adv Neur In 32.

Phaneuf, M.W. (1999). Applications of focused ion beam microscopy to materials science specimens. Micron 30, 277–288.

Pitrone, P.G., Schindelin, J., Stuyvenberg, L., Preibisch, S., Weber, M., Eliceiri, K.W., Huisken, J., and Tomancak, P. (2013). OpenSPIM: an open-access light-sheet microscopy platform. Nat Methods 10, 598–599.

Potter, C.S., Chu, H., Frey, B., Green, C., Kisseberth, N., Madden, T.J., Miller, K.L., Nahrstedt, K., Pulokas, J., Reilein, A., et al. (1999). Leginon: a system for fully automated acquisition of 1000 electron micrographs a day. Ultramicroscopy 77, 153–161.

Rigort, A., Bauerlein, F.J., Villa, E., Eibauer, M., Laugks, T., Baumeister, W., and Plitzko, J.M. (2012). Focused ion beam micromachining of eukaryotic cells for cryoelectron tomography. Proc Natl Acad Sci U S A 109, 4449–4454.

Rocklin, M. (2015). Dask: Parallel computation with blocked algorithms and task scheduling. Paper presented at: Proceedings of the 14th python in science conference.

Schaffer, M., Schaffer, B., and Ramasse, Q. (2012). Sample preparation for atomic-resolution STEM at low voltages by FIB. Ultramicroscopy 114, 62–71.

Schaffer, M., and Wagner, J. (2008). Block lift-out sample preparation for 3D experiments in a dual beam focused ion beam microscope. Microchim Acta 161, 421–425.

Schorb, M., Haberbosch, I., Hagen, W.J.H., Schwab, Y., and Mastronarde, D.N. (2019). Software tools for automated transmission electron microscopy. Nat Methods 16, 471–477.

Sofroniew, N., Lambert, T., Evans, K., Nunez-Iglesias, J., Bokota, G., Winston, P., Peña-Castellanos, G., Yamauchi, K., Bussonnier, M., Doncila Pop, D., et al. (2022). napari: a multi-dimensional image viewer for Python (Zenodo).

Suloway, C., Shi, J., Cheng, A., Pulokas, J., Carragher, B., Potter, C.S., Zheng, S.Q., Agard, D.A., and Jensen, G.J. (2009). Fully automated, sequential tilt-series acquisition with Leginon. J Struct Biol 167, 11–18.

Villinger, C., Gregorius, H., Kranz, C., Hohn, K., Munzberg, C., von Wichert, G., Mizaikoff, B., Wanner, G., and Walther, P. (2012). FIB/SEM tomography with TEM-like resolution for 3D imaging of high-pressure frozen cells. Histochem Cell Biol 138, 549–556.

Zheng, S.Q., Keszthelyi, B., Branlund, E., Lyle, J.M., Braunfeld, M.B., Sedat, J.W., and Agard, D.A. (2007). UCSF tomography: an integrated software suite for real-time electron microscopic tomographic data collection, alignment, and reconstruction. J Struct Biol 157, 138–147.

